# Entropy-based decoy generation methods for accurate FDR estimation in large-scale metabolomics annotations

**DOI:** 10.1101/2023.07.02.547371

**Authors:** Shaowei An, Miaoshan Lu, Ruimin Wang, Jinyin Wang, Cong Xie, Junjie Tong, Hengxuan Jiang, Changbin Yu

## Abstract

Large-scale metabolomics research faces challenges in accurate metabolite annotation and false discovery rate (FDR) estimation. Recent progress in addressing these challenges has leveraged experience from proteomics and inspiration from other sciences. Although the target-decoy strategy has been applied to metabolomics, generating reliable decoy libraries is difficult due to the complexity of metabolites. Additionally, continuous bioinformatic efforts are necessary to increase the utilization of growing spectra resources while reducing false identifications. Here we introduce the concept of ion entropy and present two entropy-based decoy generation methods. The assessment of public spectral databases using ion entropy validated it as a good metric for ion information content in massive metabolomics data. The decoy generation method developed based on this concept outperformed current representative decoy strategies in metabolomics and achieved the best FDR estimation performance. We analyzed 47 public metabolomics datasets using the constructed workflow to provide instructive suggestions. Finally, we present MetaPhoenix, a tool equipped with a well-constructed FDR estimation workflow that facilitates the development of accurate FDR-controlled analysis in the metabolomics field.

## Introduction

Metabolomics is the profiling of metabolites in biofluids, cells, and tissues[1]. It provides a functional readout of cellular biochemistry and is routinely applied as a tool for biomarker discovery[2]. Among all analytical techniques for untargeted metabolomics, liquid chromatography coupled to accurate tandem mass spectrometry (LC-MS/MS) has become predominant, which allows for the acquisition of thousands of metabolite signals from a single sample[3, 4]. The increased availability and innovation of instrumental techniques dramatically improve the detection and identification of metabolites[5]. To annotate and identify detected signals, tandem MS or MS/MS spectra are collected from experimental samples and compared with reference spectral libraries according to authentic standards[6, 7]. This procedure is commonly implemented by the spectrum-spectrum matching methods such as dot product similarity score[8]. However, researchers face challenges in setting appropriate scoring criteria and ascertaining the false discovery rate (FDR)[9], which can finally result in uncontrolled false positive identifications and unreliable results. The FDR is a cornerstone of the quantification of annotation quality in genomics, transcriptomics, and proteomics[10], but its immaturity in metabolomics impedes the standardization analysis in the metabolomics data processing.

To address analysis challenges in metabolite identification, the metabolomics community has leveraged experience from the proteomics field where statistical assessment and false discovery calculations for annotations are common practices[4, 11]. The representative method, FDR estimation by the target-decoy approach, has been prevalent in proteomics research for a long time but is relatively new to metabolomics due to the difficulty in generating decoy metabolomics libraries[9, 11, 12]. Despite the intrinsic complexity of metabolite fragmentation, several decoy library construction methods were proposed and validated[9, 10, 13]. The fragmentation tree-based method uses a re-rooted fragmentation tree[14] to generate decoy libraries. It was compared with the empirical Bayes approach[15] and two other decoy methods: the naïve and spectrum-based methods. The fragmentation tree-based method outperformed the other methods and was proved to provide confidence measures in large-scale metabolomics projects[9]. This is considered important progress in the target-decoy approach application in metabolomics. However, this method is limited to certain metabolomics scenarios because it can only be used on fragmentation tree-filtered spectral libraries. Another decoy generation method called the XY-Meta method was presented later and demonstrated better performance than the fragmentation tree-based method through a comprehensive comparison[13]. This method is an optimized form of the random selection method and generates forged spectra by preserving the original reference signals to simulate the presence of isomers of metabolites. Despite all these efforts from the metabolomics community, continuous methodological progress is necessary to increase the utilization of rich spectra resources while removing or reducing match redundancy[4]. This is because numerous databases are generally queried to maximize metabolome coverage and public spectral databases are growing in scale[3, 16].

In addition to obtaining inspiration from proteomics, another attempt to solve the difficulty in spectrum-spectrum matching has been made by applying information entropy theory[17] to metabolomics. Spectral entropy was introduced as a suitable measure for the total information content of an MS/MS spectrum and entropy similarity scores were proved to improve the accuracy of MS-based annotations with high robustness[18]. This successful cross-disciplines investigation demonstrates the suitability of information theory with the massive MS/MS spectral databases. However, in large-scale metabolomics analysis or downstream pathway analysis, much attention is paid to the spectrum dimension while the basic spectrum component, ion, is rarely noticed in macro statistics.

Here we introduce ion entropy as a measure for the information content of ions in MS/MS spectral libraries. We used ion entropy to assess widely-used public spectral databases including MassBank of North America (MassBank-MoNA), MassBank Europe (Massbank-Europe)[19], and GNPS[6] to reflect its applications. Based on the observed characteristics of ion entropy and previously proposed spectral entropy, we devised two entropy-based decoy spectra generation methods: the spectral entropy-based method and the ion entropy-based method. A comprehensive evaluation validated that both methods are feasible strategies to provide FDR estimation in metabolomics with high accuracy. The ion entropy-based method achieved the best FDR estimation performance when compared with other representative decoy strategies. We analyzed 47 public metabolomics datasets using our constructed workflow to validate the FDR estimation methods in real-world metabolome annotation and provided instructive suggestions. Finally, a tool named MetaPhoenix is provided to promote the development of FDR estimation in the metabolomics field.

## Results

### Ion entropy measurement on public spectral databases

Public MS/MS spectral databases typically accumulate abundant spectra recorded at multiple collision energies to cover a broad range of characteristic fragments[20]. Low collision energies mostly preserve the precursor ion, while high collision energies increase product ion abundances toward low m/z ranges and lower the precursor ion abundance (**Fig 1a**). Some ions arise in all recorded energies, while others only arise within a certain collision energy range. Product ions appearing at a wide range of collision energies might indicate that they have relatively stable substructures that are hard to dissociate.

**Fig. 1.**
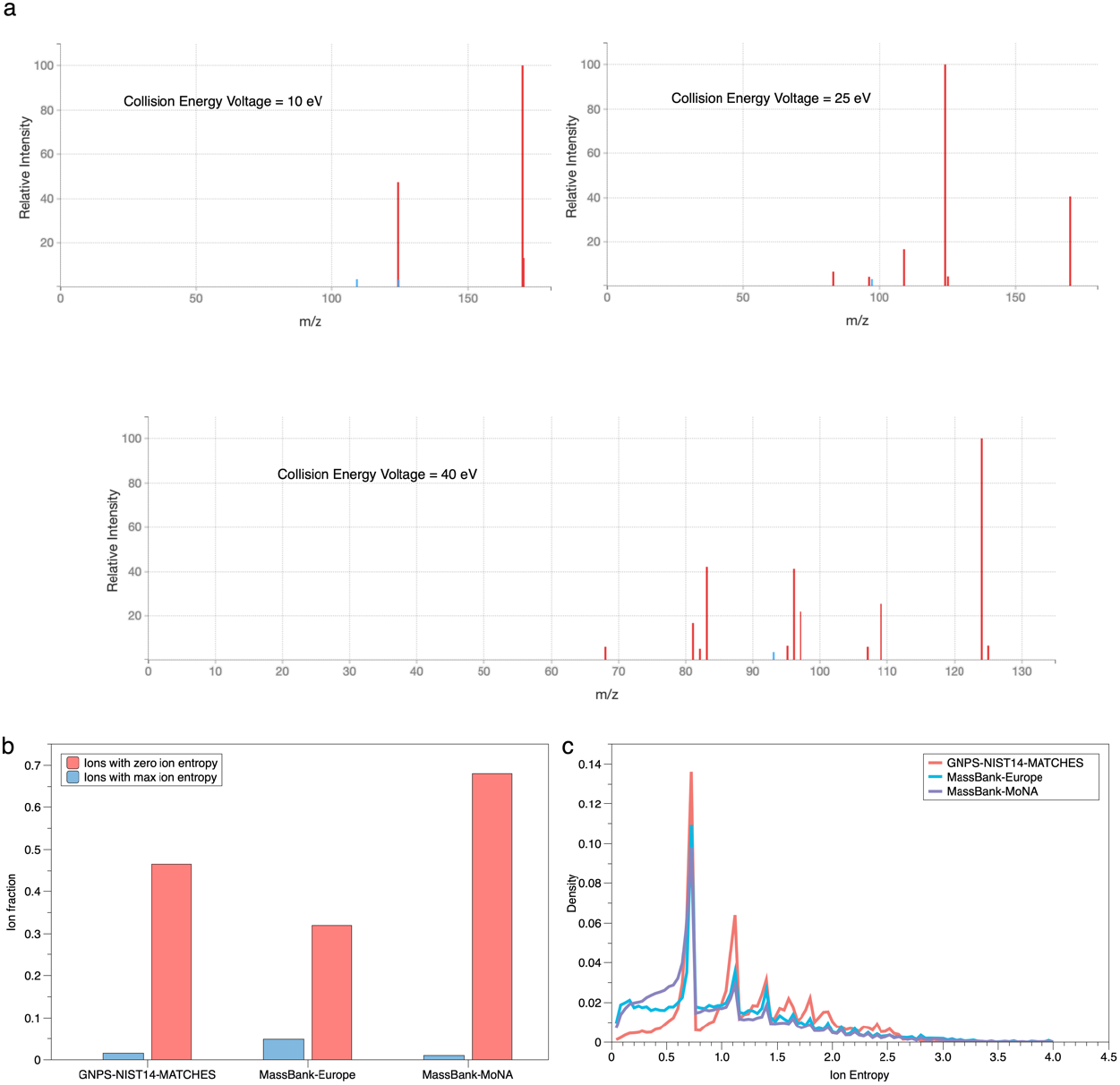
Assessment of public spectral databases using ion entropy. **a)** Spectra of 1-Methylhistidine (HMDB ID: HMDB0000001) recorded at different collision energies in HMDB. **b)** Fractions of ions with zero ion entropy and ions with max ion entropy in the GNPS-NIST14-MATCHES, MassBank-Europe, and MassBank-MoNA databases. **c)** Distribution of ion entropies in the GNPS-NIST14-MATCHES, MassBank-Europe, and MassBank-MoNA databases.

To measure the information content of ions in MS/MS spectral databases, we propose ion entropy, an application of Shannon entropy from information theory[17] into metabolomics. Entropy is originally considered a measure of disorder or uncertainty and is very suitable as an indicator for ions due to their multilevel distribution in massive spectral libraries. We assessed preprocessed public spectral databases, including GNPS-NIST14-MATCHES[6], MassBank-Europe[19], and MassBank-MoNA using ion entropy (**Fig 1**). The detailed calculation process of ion entropy is described in the methods section.

Ions with zero ion entropy and max ion entropy (value=1 after normalization) have unique characteristics. The former ions are unique in the product ion warehouse with no other reference, which may indicate a low confidence level due to the scarcity. The latter ions exist in every spectrum of a given precursor m/z value and have the same relative intensity. This kind of ion is expected to have strong stabilities during the fragmentation process because the spectra in public databases are recorded at a wide range of collision energies[21]. We calculated how much these two special ion types account for in public spectral databases (**Fig 1b**). Ions with zero ion entropy accounted for over 30%, while ions with max ion entropy accounted for no more than 5% in all these databases. These indicators have the potential to be used to measure the quality of other spectral libraries in the future. In addition, ion entropy was calculated for other ions as well, and the ion entropy distributions of these databases were illustrated (**Fig 1c**). Sharp increases in the density were observed before every *lnN* value (N represents the natural integer), and the apex density reduced as the ion entropy increased. This can be explained by the composition of the spectral databases (**Fig 1a**). Due to the ion overlap among spectra under different collision energy conditions, ion entropy will reach its relative maximal value (*Ion Entropy* = *lnN*, for N spectra) if an ion has the same relative intensity in all spectra. Moreover, ions present in fewer spectra are definitely more numerous than those present in more spectra. This theoretical conjecture is consistent with the observed phenomenon.

Overall, ion entropy can be a good metric of information content, stability, or enrichment of ions in spectral databases.

### Decoy library construction by entropy-based methods

Information entropy can be used as a key concept in the generation of decoy MS/MS spectral libraries. It is believed that the construction of decoy spectra should mimic real spectra as closely as possible but not correspond to MS/MS spectra of any true metabolites present in the sample[9]. We interpret this proposal from the perspective of information entropy theory[17] and propose that the decoy spectrum should have the same (or similar) spectral entropy as the target spectrum. Under this circumstance, they shall have similar states of chaos or disorder. Moreover, this might also suggest that the generated decoy spectrum comes from the same collision energy condition as the original target spectrum[18].

Two entropy-based methods were developed to create dependable decoy spectral libraries (**Fig 2**), called the spectral entropy-based method and the ion entropy-based method. The spectral entropy-based method randomly exchanges all ion intensities in a target spectrum to create a decoy spectrum that maintains the exact same spectral entropy as the target spectrum. The ion entropy-based method also follows this basic rule but through a more complicated process. Firstly, each ion in a target spectrum is assigned an ion entropy value and sorted by it. Then their intensities are exchanged sequentially according to their rank, meaning that ions with higher ion entropies receive intensities from the ion with lower ion entropies. Ions with zero ion entropy only exchange intensities within their own kind. This strategy is similar to the protein sequence reversal strategy[12], which is an appealing method due to its simplicity and effectiveness.

**Fig. 2.**
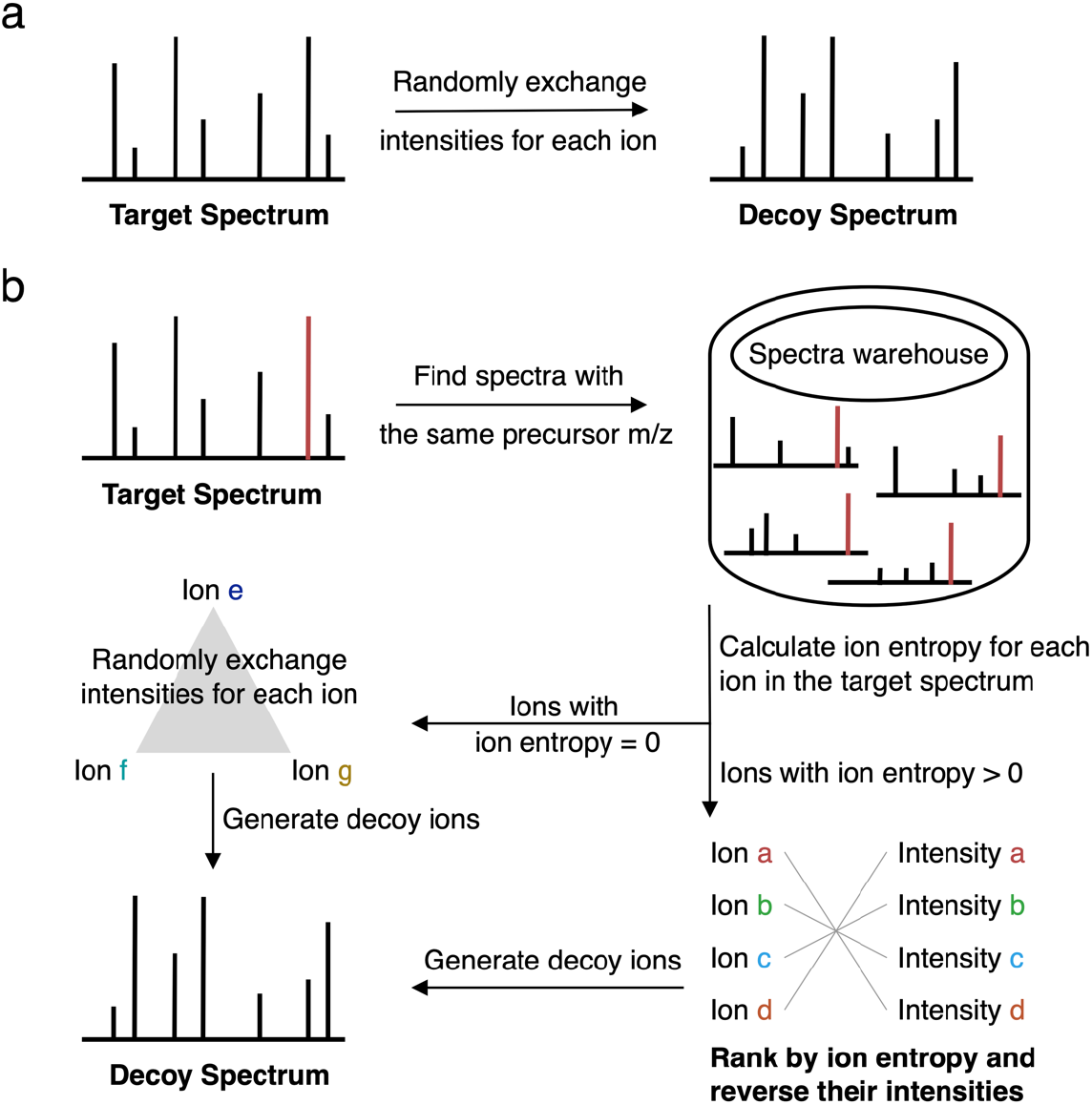
Decoy library construction schema. **a)** The spectral entropy-based decoy method. **b)** The ion entropy-based method.

### Decoy spectral database validation

Accurate control of spectral entropy during the generation of decoy spectra libraries improves our methods’ ability to mimic reference target libraries. We evaluated the spectral entropy distribution of decoy libraries generated from the GNPS[6] database using various decoy strategies (**Fig 3**). The entire GNPS library was preprocessed to satisfy the decoy generation calculation prerequisites and filtered by the fragmentation tree[22] analysis to be adopted for the fragmentation tree-based method. It was observed that all other methods broke the original spectral entropy distribution of the target library except for the entropy-based methods. The naive and XY-Meta methods caused significant shifts in the distribution shape towards lower spectral entropies. The fragmentation tree-based method was basically similar to the original distribution for the filtered library while the entropy-based methods maintained perfectly consistent distributions with the target library both for the filtered or unfiltered conditions. Surprisingly, although generating decoy libraries using a mechanism completely irrelevant to entropy, the fragmentation tree-based method obtained similar results to our entropy-based methods.

**Fig. 3.**
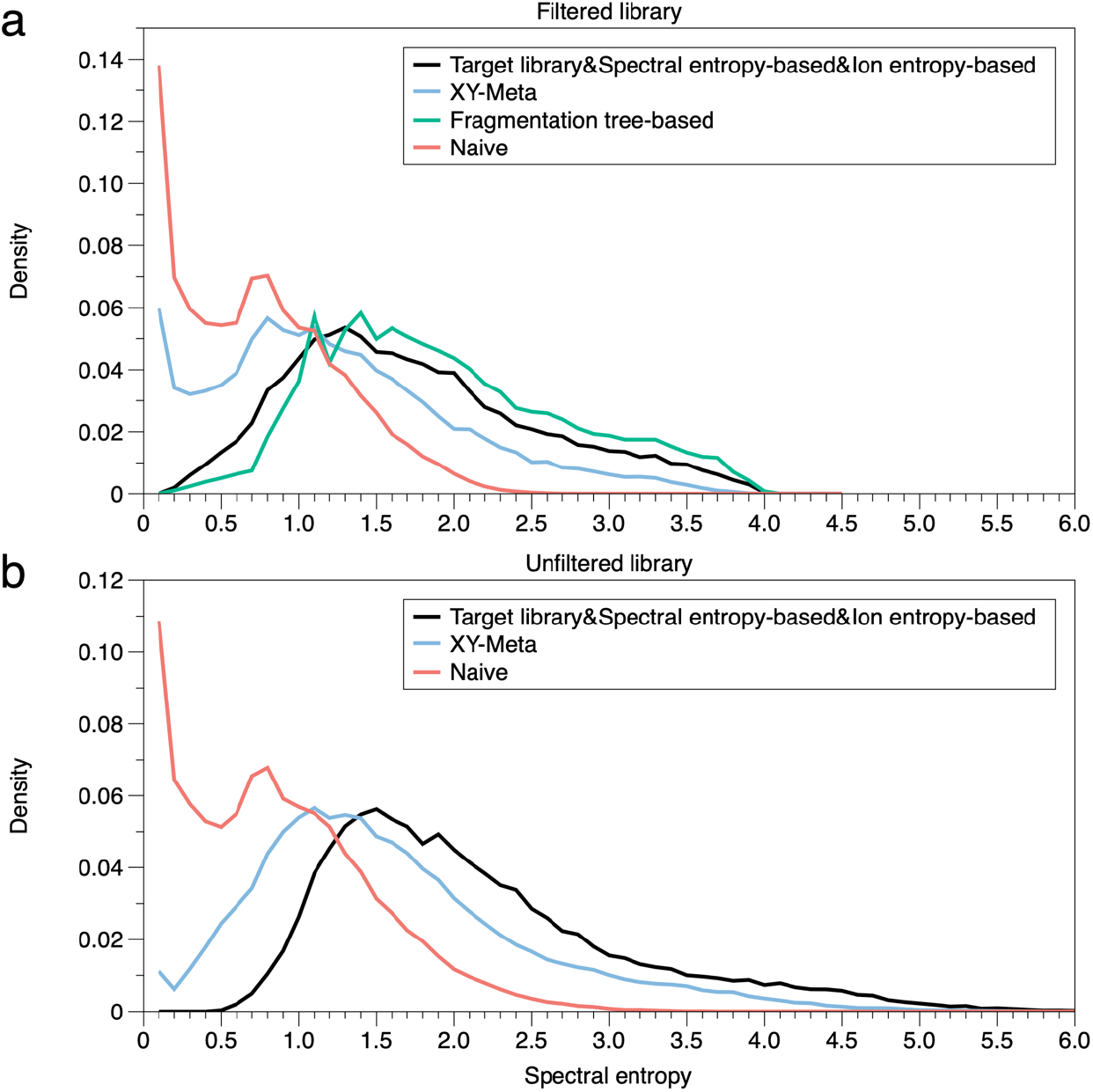
Spectral entropy distributions of decoy libraries generated by different decoy strategies. **a)** Using the fragmentation tree filtered GNPS library and **b)** unfiltered library to generate decoy libraries.

P-value estimation[23] was used to further validate the entropy-based methods and assess the decoy database quality. We conducted a “separated target-decoy search” (STDS)[14] analysis using the MassBank-MoNA library as the query library and the GNPS library as the target library. The two entropy-based strategies were used to generate decoy libraries from the GNPS library. False positive identifications can be calculated using the first part of InChIKeys, and we can investigate if p-values of these false hits estimated by our methods are uniformly distributed[23]. Mostly uniform distributions of p-values were observed both for the spectral entropy-based method and the ion entropy-based method (**Fig 4**), which agrees with the distribution of p-values under the null hypothesis. The p-value of a spectrum match is defined as the probability to randomly draw a result of this or better quality under the null hypothesis for which a spectrum has been randomly generated[9]. Our decoy strategies were validated as good decoy generation methods by both spectral entropy distributions and p-value estimation.

**Fig. 4.**
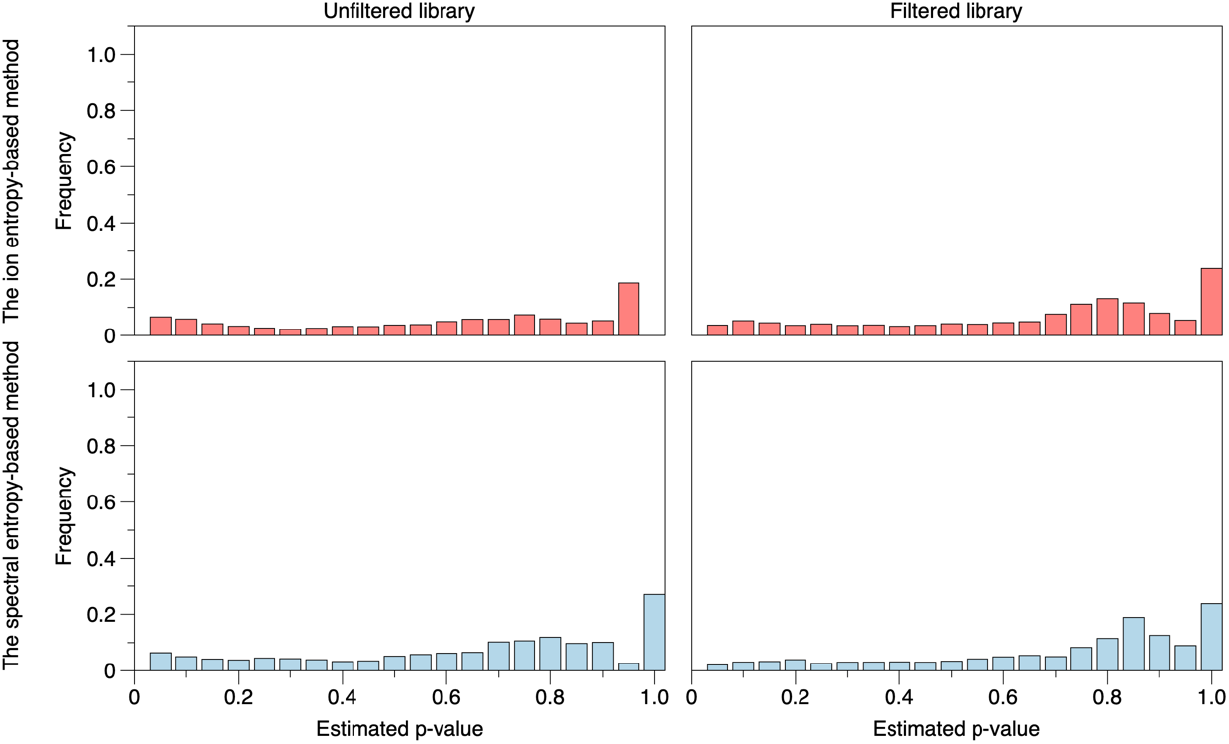
Distributions of p-values for false identifications. The MassBank-MoNA query spectra were used to search the GNPS library and decoy libraries generated by the entropy-based methods. The fragmentation tree filtered and unfiltered libraries were both assessed.

### Accurate FDR estimation and performance comparison

Our methods were assessed by observing how well the estimated FDR mimics the real FDR. Three other decoy generation methods were used for the performance comparison under the same condition. The naive method[9] is a simple random method that generates decoy spectra by randomly picking ions from the whole target library until the decoy spectrum mimics the corresponding target spectrum. The fragmentation tree-based method[9] uses a re-rooted fragmentation tree[22] to generate decoy libraries while the XY-Meta method[13] is an improved version of the random selection method and achieved better performance than the fragmentation tree-based method on certain datasets. We chose the MassBank-MoNA library and the GNPS library as the query library and the target library. Decoy libraries with and without the fragmentation tree filter were generated since the fragmentation tree-based method can only be used on fragmentation tree-filtered databases.

Both tested situations showed that the entropy-based methods performed well (**Fig 5**). The spectral entropy-based method showed good FDR estimation especially at the low FDR range (0-0.1) while the ion entropy-based method traced the expected curve best and accurately evaluated the true FDR almost in the entire range (0-0.5). The result demonstrates that generating decoy libraries by accurate entropy control is effective for both filtered and unfiltered databases. Overall, the ion entropy-based method performed better than other existing methods while the spectral entropy-based method could also be used for FDR estimation under certain circumstances. It should be noted that estimating FDR based on unfiltered target libraries might be preferred in most scenarios, in which case our ion entropy-based method demonstrates the best applicability due to its accurate prediction of FDR.

**Fig. 5.**
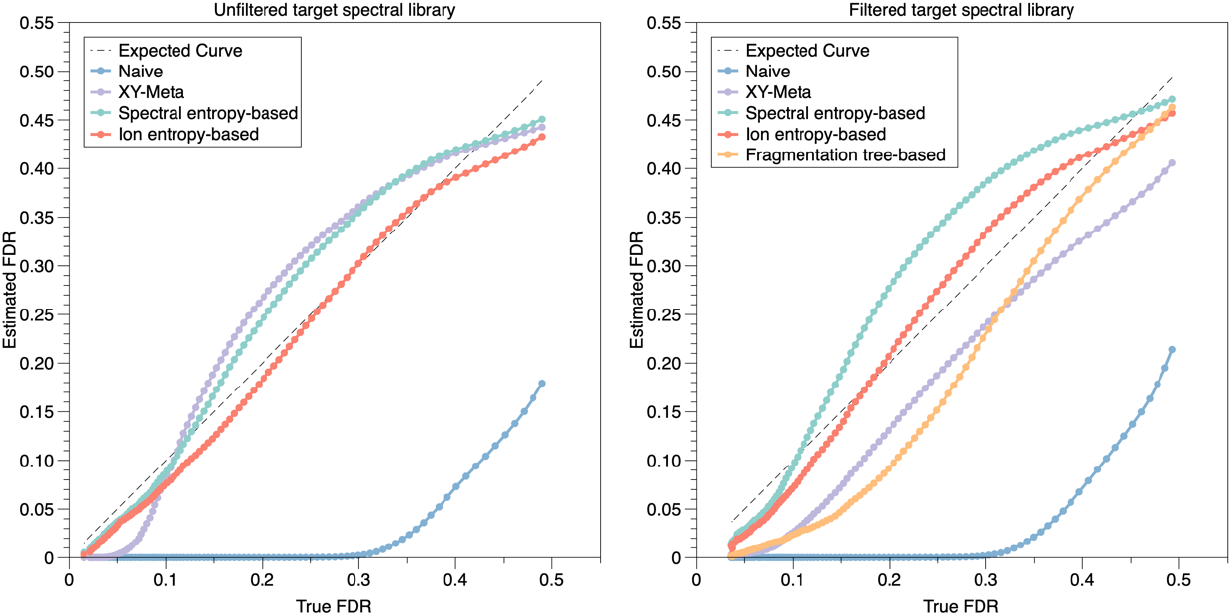
FDR estimation performance comparison. The MassBank-MoNA query spectra were used to search the GNPS database and its decoy libraries to calculate the estimated FDR. The fragmentation tree filtered (right) and unfiltered libraries (left) were tested.

### Public metabolomics datasets validation

To demonstrate how our methods can be used in real metabolomics datasets, we conducted FDR estimation analysis on 47 public datasets based on the ion entropy-based decoy strategy and observed the spectrum identification fractions (**Fig 6**). The whole GNPS library was used as the reference library after preprocessing. 56480 spectra were reserved for the analysis and decoy library generation. We chose 0.01, 0.05, and 0.1 as the FDR control thresholds. In the threshold of 0.01, the identification fractions for most datasets were less than 10%. When it comes to 0.05, the identification fractions increased remarkably while at the threshold of FDR<0.1, the spectra utilization became better but remains at a limited level. Since the ion entropy-based method is considered to estimate the FDR accurately (**Fig 5**), this result hints that using accurate FDR control to conduct metabolite identification is very necessary due to the large potential proportion of false positives.

**Fig. 6.**
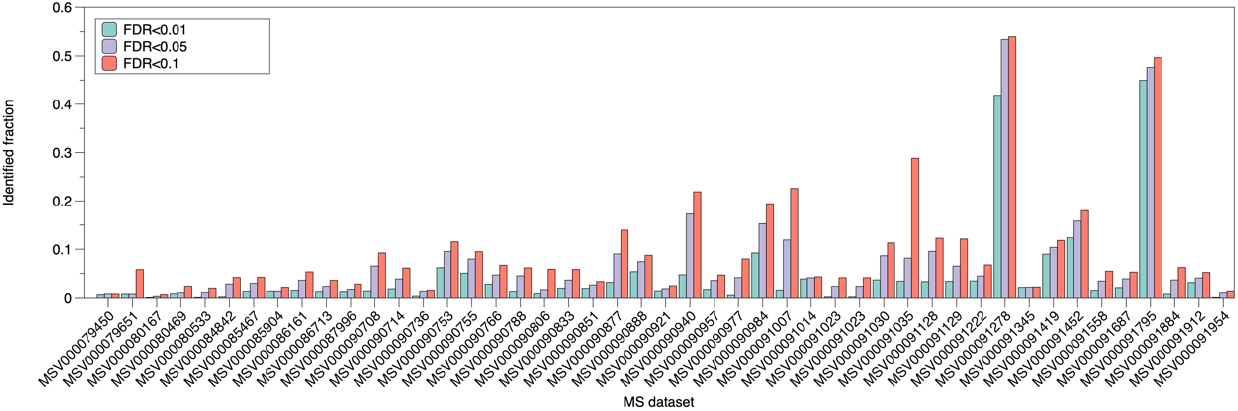
Metabolite identification on public datasets. The ion entropy-based decoy method was used for accurate FDR estimation.

**Fig. 7.**
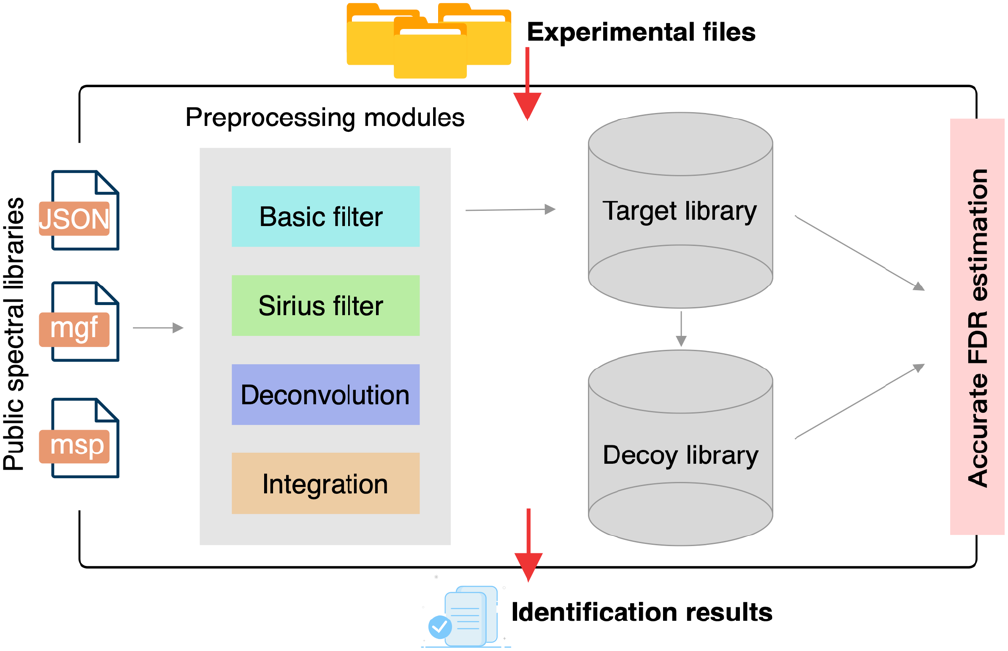
Schema of MetaPhoenix workflow.

### Software with well-constructed target-decoy analysis workflow

Our methods and workflow have been packaged into an easy-to-use tool called MetaPhoenix for researchers to conduct relative standard FDR estimation analysis considering the prevalence of the target-decoy approach in proteomics but the immature in metabolomics. This tool takes JSON, mgf or msp format files as input stream and txt or excel format files as output.

Multiple function modules are provided for preprocessing of reference target libraries while supporting linkage to other software such as SIRIUS[24]. The Spectra deconvolution function was developed to satisfy common analysis needs and spectra can also be integrated according to certain items like SMILES or InChIKeys to eliminate duplication and redundancy. We expect this tool can resolve the analysis difficulty in generating decoy libraries and accurate FDR estimation in metabolomics.

## Discussion

Ion entropy was introduced as a metric for measuring ion information in ever-increasing tandem MS/MS spectra in databases. Unlike the relatively broad definition of spectral entropy, only product ions with the same precursor m/z were considered by the calculation process. This reduces the impact from other unrelated ions and better reflects the characteristics of certain product ions. In LC-MS/MS-based metabolomics, ion information is usually noticed and used when researchers need to annotate m/z values in a single spectrum or search compounds for a specific m/z value in spectral databases. However, it is given little attention in statistics for metabolomics-based biological discoveries. As we gain more authentic spectra from labs, data mining on these accumulated spectra may lead to effective discoveries. If we can find the relationship between ion entropy and the molecular collision process based on a large scale of collected spectra, it will be possible to build simulation methods that generate reliable in silico spectra or measure the quality of existing spectral libraries.

The use of ion entropy in generating decoy libraries is similar to the peptide reversal strategy in proteomics. One of the main challenges in generating metabolomics decoy spectra is that small molecules are structurally diverse and therefore difficult to shuffle or reverse[9]. The ion entropy-based method addresses this challenge by providing a quantifiable characteristic to ions based on the evaluation of existing spectral databases. This indicator allows diverse ions to be distinguished instead of relying solely on m/z values.

Diverse ions can now be shuffled according to certain patterns like how we shuffle the 20 amino acids in proteomics. It should be noted that multiple factors can affect ion entropy, such as ion abundance, stability, isomerization, collision energy distribution, and quality of MS/MS spectral libraries. Low-quality spectral libraries may have adverse effects on the accurate calculation of ion entropy due to uncontrolled spectrum repetition or narrow distribution of collision energies. Considering the dependency of ion entropy calculation on spectral library quality, we suggest using the entropy-based decoy generation methods on authentic huge spectral libraries for accurate FDR estimation.

Using spectral entropy similarity as the spectrum-spectrum matching criterion in our workflow has several advantages. Firstly, it outperforms all classic algorithms like dot product similarity and minimizes the impact of wrong spectral matches on the final FDR estimation[18]. Additionally, there is no need to set a minimum matching ion count parameter, which avoids confusion for researchers. Finally, it has a dynamic exponential weight function with subsequent normalization applied to spectra with low spectral entropy that helps distinguish highly similar structures. These advantages led us to adopt it as the suggested similarity measure instead of the commonly used dot product similarity.

In summary, our work extended the application of information entropy in metabolomics by proposing a quantity to measure the information content of ions in spectral libraries. Based on this new concept, we developed novel decoy spectral library generation methods that outperform current representative metabolomics decoy strategies. Our methods achieved more accurate FDR estimation on large-scale metabolomics analysis through comprehensive validation. MetaPhoenix is provided to standardize the FDR estimation analysis in metabolomics.

## Methods

### Ion Entropy Calculation

To calculate ion entropy for an ion in an MS/MS spectrum, firstly all the spectra that have the same precursor m/z with this spectrum were collected from the reference database. Then each spectrum was normalized by dividing their ion intensities by the precursor ion intensity. These normalized ions made up an ion warehouse. Ions within the range of 10 p.p.m from the target ion m/z were picked out from the warehouse and their intensities were adjusted to have sum=1 by dividing their sum intensity. Ion Entropy (IE) can then be calculated from these ion intensities *I*_*p*_ by the equation:

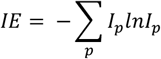

Normalized ion entropy can be calculated by the equation:

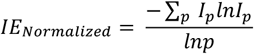

### Spectral library preprocessing

All the query and target spectral libraries were preprocessed by the following steps to ensure maximal homogeneity. Spectra were removed if they have:

1. key information (m/z values, intensity values, precursor m/z, ion mode, InChIKey) missing.
2. less than 5 peaks with relative intensity above 2%.
3. no ion within 10 p.p.m range of the precursor m/z.
4. precursor m/z (or exact mass) over 1000.
5. other spectrum types rather than MS2 type.
6. negative or unknown ion mode.

Besides, ions with relative intensity <1% were treated as noise signals and abandoned.

### Fragmentation tree-based processing

The fragmentation tree-based decoy library generation and noise filtering were conducted by SIRIUS 5.6[24]. The running parameters were set as default parameters.

### Spectral matching function

We implemented the entropy similarity score function and used it for spectral similarity measurement. The similarity between spectrum A and spectrum B was calculated after they were mixed to get spectrum AB by the equation:

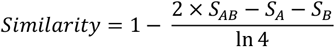

Details about this score function could be found in its origin publication[18].

### FDR calculation

We used STDS strategy[14] for the FDR estimation under which circumstances all the library hits above a specific threshold would be reported. The corresponding FDR is the total number of above-threshold decoy library hits (N_D_) divided by the number of above-threshold target library hits (N_T_).

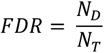

## Data availability

Public spectral databases MassBank-MoNA (https://mona.fiehnlab.ucdavis.edu), MassBank-Europe (https://massbank.eu/MassBank) and GNPS (https://gnps.ucsd.edu) were used for this study. Real metabolomics datasets used were downloaded from GNPS.

## Code availability

https://github.com/anshaowei/MetaPhoenix

